# Passive Exposure Sparsifies Neural Activity in the Primary Visual Cortex

**DOI:** 10.1101/2021.11.18.469160

**Authors:** Jan Homann, Hyewon Kim, David W. Tank, Michael J. Berry

**Affiliations:** Princeton Neuroscience Institute, Princeton University, Princeton, NJ 08544

## Abstract

A notable feature of neural activity is sparseness – namely, that only a small fraction of neurons in a local circuit have high activity at any moment. Not only is sparse neural activity observed experimentally in most areas of the brain, but sparseness has been proposed as an optimization or design principle for neural circuits. Sparseness can increase the energy efficiency of the neural code as well as allow for beneficial computations to be carried out. But how does the brain achieve sparseness? Here, we found that when neurons in the primary visual cortex were passively exposed to a set of images over several days, neural responses became more sparse. Sparsification was driven by a decrease in the response of neurons with low or moderate activity, while highly active neurons retained similar responses. We also observed a net decorrelation of neural activity. These changes sculpt neural activity for greater coding efficiency.

## Introduction

One of the striking properties of brain activity is the fact that very few neurons are active at any moment. This fact was not always appreciated in early recordings with extracellular electrodes, because of the strong bias that technique has for sampling highly active neurons (Brecht and Sakmann, 2002; DeWeese et al., 2003; Harris et al., 2000; Shoham et al., 2006). However, these earlier methods did show that neural activity was sparser when stimulated outside of the classical receptive field (Haider et al., 2010; Vinje and Gallant, 2000) or when compared to the activity estimated from classical receptive field models (Weliky et al., 2003). Using techniques with much lower bias, such as cell-attached patch recordings, silicon probes, and two-photon calcium imaging, neural activity has been found to be highly sparse in many cortical areas (Froudarakis et al., 2014; Hromadka et al., 2008; Kampa et al., 2011; O’Connor et al., 2010; Shoham et al., 2006; Yen et al., 2007; Yoshida and Ohki, 2020) as well as many subcortical structures (Berry et al., 1997; Butts et al., 2007; Hahnloser et al., 2002; Ito et al., 2008). A closely related phenomenon is the observation that the distribution of firing rates in many brain areas follows a log-normal distribution with high enough skew to be sparse (Buzsaki and Mizuseki, 2014).

Early insight into the importance of sparse neural activity came from a study showing that a sparse neural code that represents natural images with low error uses Gabor-like receptive fields, as found in the primary visual cortex (Olshausen and Field, 1996a). Sparseness can similarly account for the receptive field structure of auditory nerve fibers (Smith and Lewicki, 2006). More generally, sparse codes are well-matched to the statistics of natural sensory stimuli (Field, 1987; Hyvärinen et al., 2009; Olshausen and Field, 1996b; Simoncelli and Olshausen, 2001). In fact, the power-law distribution of population activity patterns found in both retina and cortex (Tkacik et al., 2015; Yu et al., 2013) closely matches the distribution of image patches in natural scenes (Stephens et al., 2013). Another advantage of a sparse code is energy efficiency (Balasubramanian et al., 2001; Laughlin, 2001; Levy and Baxter, 1996), as neural activity requires substantial metabolic energy (Attwell and Laughlin, 2001). An analysis of the energy budget in the cerebral cortex suggests that neurons may, in fact, be constrained to low average activity (Lennie, 2003). Sparse codes can also be easier for subsequent neural circuits to read out (Barak et al., 2013; Olshausen and Field, 2004), enhance learning (Ito et al., 2008; Lin et al., 2014; Schweighofer et al., 2001), and increase the capacity of associative memories (Baum et al., 1988; Treves and Rolls, 1991).

Given these many advantages, it is natural to ask how neural circuits might achieve sparse codes. Local circuit mechanisms, such as lateral inhibition (Theunissen, 2003), adaption (Betkiewicz et al., 2020), thresholding (Paiton et al., 2020), and coincidence detection linked to input synchrony (Perez-Orive et al., 2002) can all create sparse neural codes. However, all of these mechanisms require tuning to produce useful levels of sparseness; in particular, overly sparse codes represent very little information (Laughlin, 2001). How neural circuits choose beneficial sparseness is not known. One possibility is that evolution has shaped beneficial levels of sparseness. Early in development, the visual cortex exhibits waves of correlated activity (Rochefort et al., 2009), which then refine to sparser activity (Golshani et al., 2009). However, later in development sparseness appears to decrease (Berkes et al., 2009). More generally, local neural circuits in the neocortex and many other brain areas exhibit high levels of synaptic plasticity throughout adult life (Feldman, 2009; Ribic, 2020) that interact to maintain homeostasis (Turrigiano, 2011). Models that combine Hebbian, anti-Hebbian, and homeostatic plasticity have been shown to create sparse neural codes (Foldiak, 1990; Zylberberg et al., 2011).

While sensory tuning in the adult has been thought to be relatively hard-wired, recent experiments demonstrate that repeated exposure to visual stimuli can increase the local field potential (Frenkel et al., 2006) or the intrinsic imaging signal (Kaneko et al., 2017) evoked by those stimuli.

However, at the neuronal level, the picture is mixed, with some studies reporting that this paradigm induces a net decrease in activity (Henschke et al., 2020; Kim et al., 2019b; Makino and Komiyama, 2015) and another study reporting a net increase (Kaneko et al., 2017). Extreme interventions, such as dark exposure (Solarana et al., 2019) and associative fear conditioning (Gdalyahu et al., 2012), can also increase sparseness, but it is not known whether passive visual exposure can have such an effect.

Here, we study how the sparseness of the neural code changes during repeated visual exposure to a set of visual images. The animals did not perform tasks or receive rewards, so only unsupervised learning mechanisms were at play. We recorded from layer 2/3 neurons in the primary visual cortex of awake, head-fixed mice using two-photon calcium imaging. We found that the sparseness of the population neural code, in fact, increased over six days of visual exposure. This effect was not caused by increased activity of the most strongly responding neurons. Instead, it emerged from a decrease in visual responses of the majority of weakly active neurons. We also found that the correlation between pairs of neurons decreased during visual exposure. Both effects resulted, in part, from a sharpening of the tuning curve of neurons, which also increased the lifetime sparseness of individual neurons. Together, these changes suggest that multiple plasticity mechanisms interact to sculpt and control the sparseness of the neural code in the adult cortex.

## Results

In order to test if the sparseness of the population code changes with visual exposure, we presented repeated images to four awake mice for six consecutive days. Images were projected on a toroidal screen surrounding the mouse. During presentation, mice were head fixed and placed on an air suspended Styrofoam ball that allowed them to run freely (Fig. 1A). Neural activity was recorded in layer 2/3 of V1 with a two-photon microscope through a glass covered cranial window (Fig. 1B). During the recording, animals did not perform any task. The microscope’s field-of-view was ∼400 µm x 400 µm containing on the order of 100 neurons per mouse. Slight day-to-day shifts in the z-plane of the field-of-view as well as intrinsic brain motion made some neurons drop out of view on some days. Roughly 50% of all recorded neurons could be tracked across all 6 days. Only those neurons were used for analysis. In order to reduce bias in the overall activity of neurons, we identified ROIs by hand, based on time-averaged still frames. ROIs were identified independently for each day and then matched across days (see Supp. Figure 3). Raw fluorescence values were stable within a recording session and were within a few percent across days, indicating that bleaching was not significant under our conditions (Supp. Figure 4).

**Figure 1.**
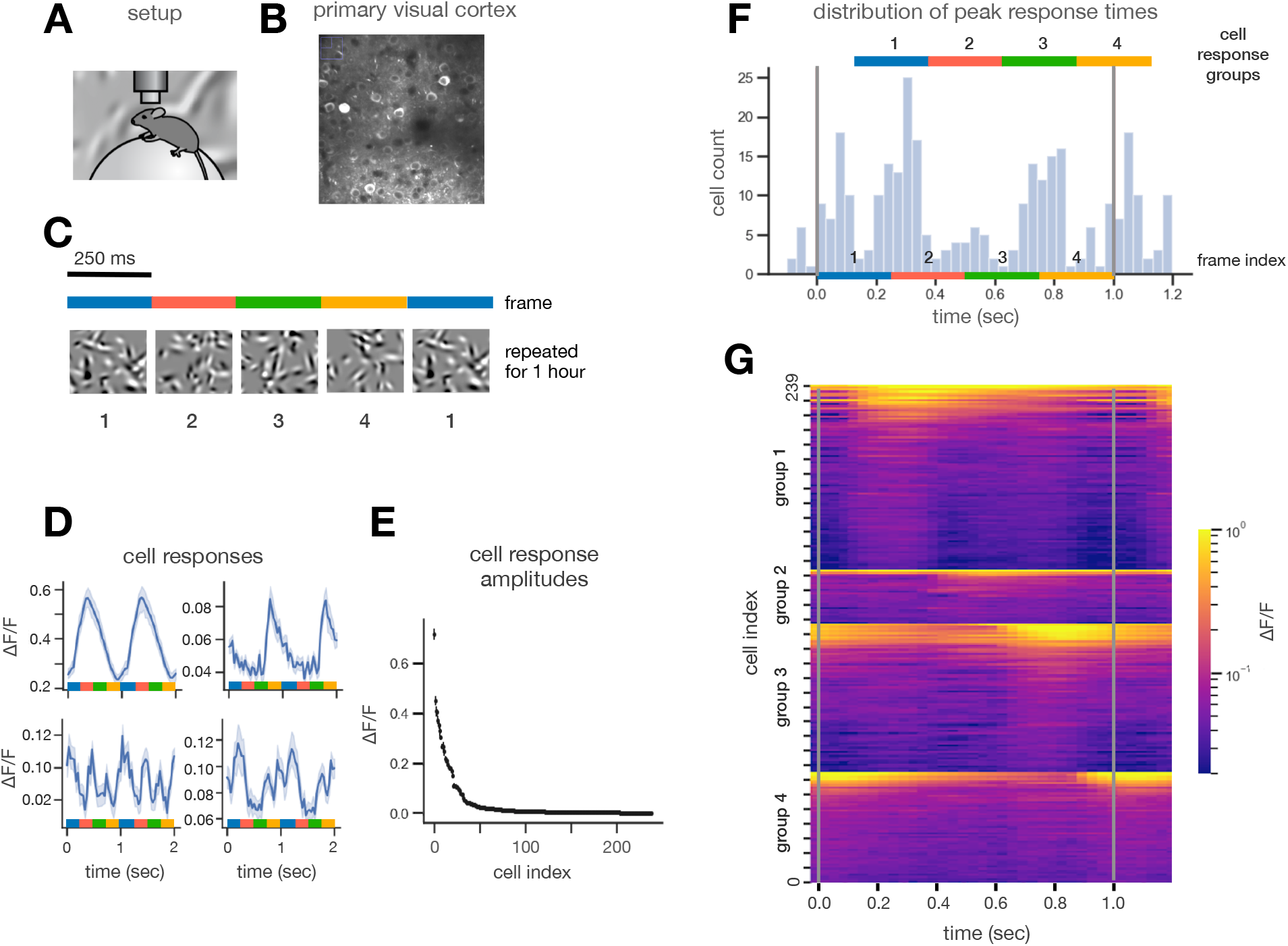
Experimental Design. **A**. *Setup*: A mouse was placed on an air suspended styrofoam ball and head fixed. Images were projected on a spherical projection screen while a two-photon microscope recorded fluorescent activity in pyramidal cells in layer 2/3 of V1 through a cranial window. **B**. Part of a typical field of view showing neurons expressing GCaMP6f. Around 50 cells per animal (N = 4) were captured across 6 days. **C**. *Stimulus design:* A 4 image sequence was presented to the mouse for 250 ms per image consisting of randomly chosen Gabors. The sequence was looped for about one hour per day for 6 days. **D**. *Cell PSTH:* Responses from 4 cells on day 1 averaged over all repeats of the sequence. **E**. *Amplitude distribution of cell responses:* Rank-ordered cell response amplitudes (ΔF/F fluctuations: defined as standard deviation of ΔF/F) for all cells on day 1. The activity of cells was highly skewed. **F**. *Histogram of peak response times:* Neurons fell into 4 groups corresponding to their preferred image. **G**. *Heatmap plot showing all cells:* X-axis: time, y-axis: cell index, color: response amplitude. Cells were first clustered by preferred image and then sorted by peak amplitude. Only a few cells responded strongly to a given frame.

The stimulus consisted of four images shown for 250 ms each in a repeated sequence that lasted for roughly one hour (Fig. 1C). Each image consisted of a superposition of randomly chosen Gabor functions, which have been shown to strongly drive neurons in the primary visual cortex (Homann et al., 2017; Kim et al., 2019a). Individual Gabor functions had sizes and spatial frequencies resembling the parameters of receptive fields measured in mouse V1 [(Niell and Stryker, 2008); see Methods]. All images were unique, yet had the same statistics, so that there were no overall changes in luminance or contrast.

We averaged neural responses across all trials on a given day (Fig. 1D), as activity on individual trials was generally intermittent (de Vries et al., 2020). Due to the large number of trials (roughly 3600), the standard error was small enough to resolve individual peaks in activity locked to the stimulus for most neurons. Responses of strongly responding cells generally had one dominant peak (Fig. 1D *top*) locked to one preferred image. Responses of weakly responding cells often had several peaks, locked to several images (Fig. 1D *bottom*).

We defined the response amplitude of individual neurons as the standard deviation of their event-triggered stimulus average (ETA; see Methods). We made this choice because many neurons had sustained baseline responses that were potentially not driven by the stimulus. Furthermore, the baseline activity changed from day to day, in part due to slight changes in the depth of our two-photon imaging. Therefore, in order to avoid excess variation in our measurement of neural responses, we excluded baseline activity. We chose the standard deviation of the response as a measure of response amplitude rather than peak-to-peak amplitude, because standard deviation is more robust to noise and because it includes activity triggered by all images.

The distribution of response amplitudes was highly skewed, with a small number of neurons exhibiting large responses and the vast majority showing small responses (Fig. 1E). This pattern of activity has often been described as “sparse” (de Vries et al., 2020; Froudarakis et al., 2014; Vinje and Gallant, 2000; Willmore and Tolhurst, 2001). One popular definition of sparseness compares the average of each neuron’s squared activity versus the average neural activity [see Methods] (Vinje and Gallant, 2000). If all neurons have identical activity, this population sparseness measure, S, is zero. If only one neuron is active and all other neurons are silent, then the population sparseness S is one. For our data, we found that the population sparseness was 0.82, similar to a large-scale survey of V1 using different choices of visual stimuli (de Vries et al., 2020).

In order to better understand how neurons responded to the stimulus, we binned the time of each neuron’s peak activity in a histogram (Fig. 1F). The vertical lines in the histogram demarcate one sequence of the four images. The color strip below the histogram indicates the time periods in which the four different images were shown. This histogram exhibits bumps that are separated by ∼250 ms. Although it is not essential to our subsequent analysis, we interpret each bump as the set of neurons that respond maximally to the same image. These histogram bumps were not evenly populated, as some images happened to evoke peak activity in more neurons than others.

We then separated the neurons into four classes, each containing the set of neurons that preferred the same image. To do this, we divided the histogram into four equal time periods of 250 ms (Fig. 1F *top*; color strip). The beginning of each time period aligned with the onset of an image, plus a response lag of 133 ms.

To visualize all neural responses, we plotted neural activity as a heatmap. In this heatmap, we grouped neurons by their preferred image and within each group by activity (Fig. 1G). Vertical stripes were visible, each corresponding to the set of neurons whose peak response was to the same image. Only a few neurons had high activity within each group, showing that population neural activity was sparse for every image.

### Changes in Population Activity Patterns

We next examined how neural activity changed across days of repeated stimulation by the same set of images. We observed many patterns of changes in response across days. Some neurons exhibited a response amplitude that decreased systematically from day-to-day with relative responses to all images remaining similar (Fig. 2A *top*). Some neurons that had a strong response to one image showed little decrease in amplitude. However, many of these neurons had a small secondary response (seen as a shoulder on the main peak) that decreased significantly across days (Fig. 2A *middle*). Finally, other neurons had lower activity with multiple peaks in their response. Often, the smaller peak decreased more in amplitude than the primary peak (Fig. 2A *bottom*).

**Figure 2.**
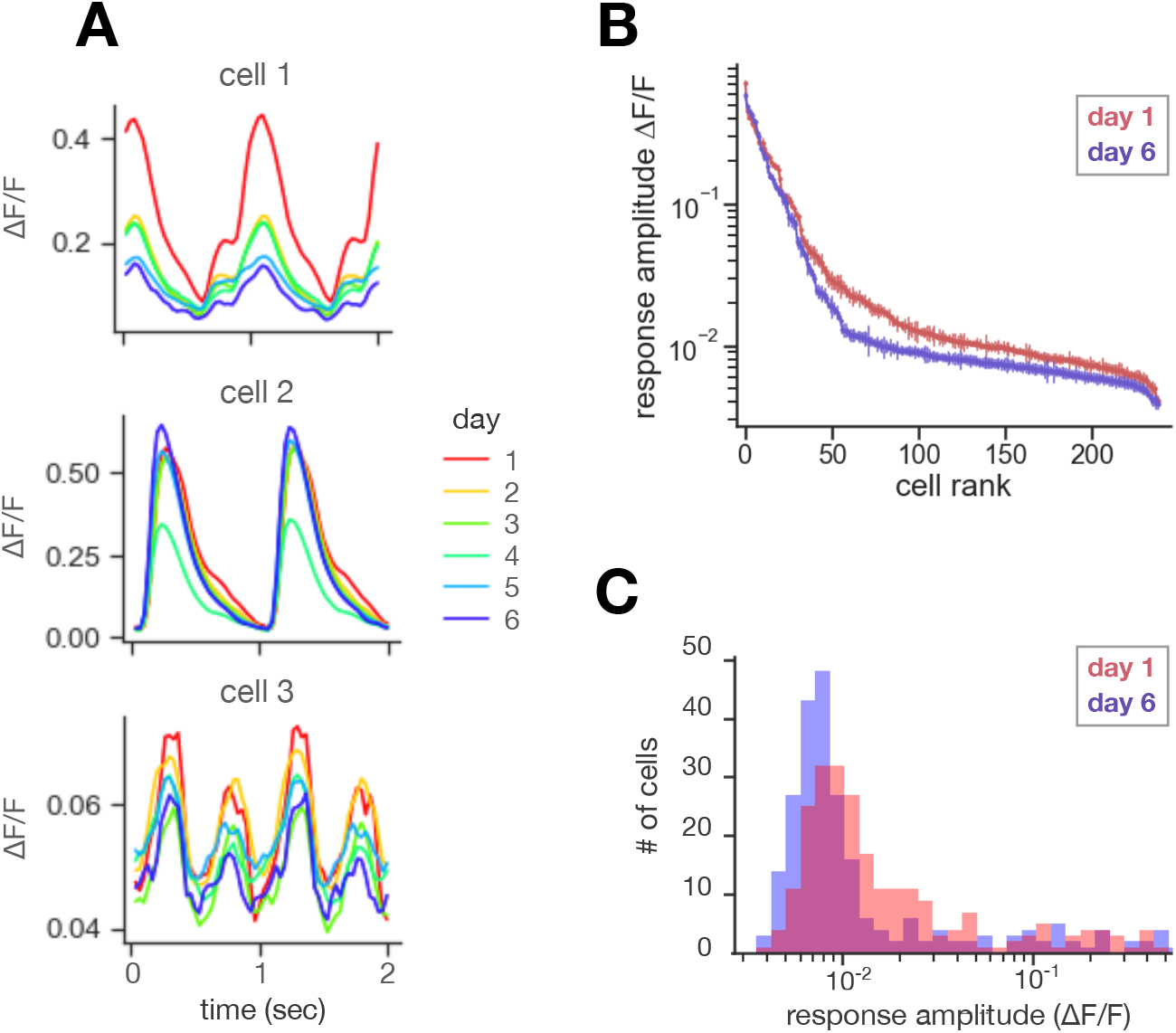
Population Sparseness. **A**. *Change in responses across days for 3 cells:* Cell 1 decreased its amplitude but the overall shape of the response stayed similar. Cell 2 had an amplitude that is roughly stable but a small secondary response hump disappeared. In cell 3 primary and secondary response changed differentially, with the secondary response decreasing more than the primary response. **B**. *Empirical cumulative distribution of response amplitudes on day 1 and day 6:* The response amplitude of most cells decreased and the distribution of responses became more skewed from day 1 to day 6. Note that the y-scale is logarithmic. **C**. *Histogram of response amplitudes*. Again, it can be seen that the average response went down and the distribution became more skewed. X-scale is logarithmic.

To compare how the pattern of activity across the population changed, we compared the rank-ordered responses on day 1 and day 6 (Fig. 2B). Because response amplitudes were highly skewed, we used a log scale. The comparison showed that a small subset of the most active neurons (roughly the top 25 by rank) had a very similar distribution of responses across days. In contrast, the majority of less active neurons (roughly the bottom 200 by rank) showed substantially smaller responses on day 6 than day 1.

To provide additional visualization, we re-plotted the same data as a histogram on a log scale for day 1 versus day 6 (Fig. 2C). This display showed a small tail of larger responses that remained similar across days, along with a roughly log-normal distribution of lower activity that decreased across days.

All together, these results showed that a small subset of the most active neurons accounted for a larger share of total activity after six days of passive viewing of the same images. The mechanism underlying this change was not an increase in activity of active neurons, but instead a decrease in activity of the other neurons. These qualitative changes also suggest an increase in the sparseness of the population code.

### Quantifying Population Sparseness

In order to quantify these observations, we computed several measures to characterize the asymmetry of population activity and followed their values across days of exposure to the same visual stimuli. First, we computed the population sparseness, *S*. There was a general trend towards higher values of population sparseness across days of exposure (Fig. 3A). In order to estimate the significance of this trend across days, we found the best linear fit and calculated the significance of this slope being greater than zero. This trend was significant (p = 0.018).

**Figure 3.**
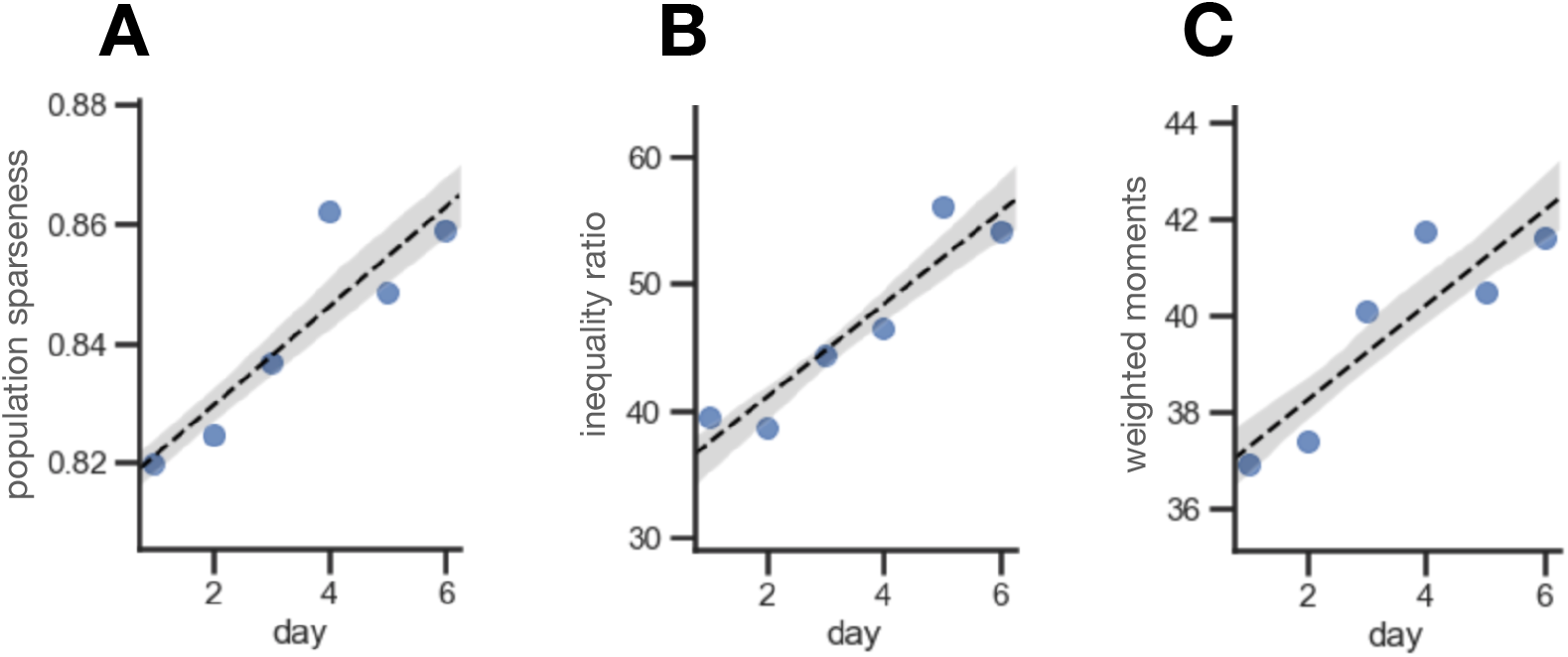
Quantifying Sparseness. **A**. *Population sparseness, S* (circles), plotted as a function of days of visual exposure; the slope of the linear fit (dotted line) was significantly different from 0 (p = 0.018). **B**. *Inequality ratio, I* (circles), plotted as a function of days of visual exposure; again, the slope of the linear fit (dotted line) was significantly different from 0 (p = 0.006). **C**. *Weighted higher-order moments, M*, plotted as a function of days of visual exposure; again, the slope of the linear fit (dotted line) was significantly different from 0 (p = 0.02). **A-C:** Gray error bands indicate the 68% confidence region of the linear fit, determined through bootstrap resampling.

Next, we computed the inequality ratio, *I*, which is defined as the average activity of the 10% most responsive neurons divided by the average activity of the 50% least responsive neurons (see Methods). This measure directly captures the qualitative changes in the pattern of population activity that we observed, and its value is easily interpreted. However, another motivation for our specific choice of the definition of *I* is that this same measure has been studied in the economics literature, where it quantifies the degree of income inequality in human populations (Piketty and Goldhammer, 2014). Choosing to compute the exact same measure opens up the possibility of further connections emerging between these seemingly disparate fields.

The overall values of the inequality ratio were quite high, ranging from 40 up to 55. These values underscore the extremely long tails present in the pattern of population activity. In addition, there was a clear and significant trend, in which *I* increased across days of passive exposure (p = 0.006).

Finally, we also computed the first four statistical moments of the distribution of population activity. For greater robustness, we calculated these statistical moments in the distribution of log activity (Fig. 2C). Both the mean and median decreased across days (Supp. Fig. 1; mean, p = 0.001; median p = 0.0004). At the same time, the higher moments increased across days (standard deviation: p = 0.33, skew: p = 0.088, kurtosis: p = 0.2). Because of the moderate statistical significance of these trends, we constructed a composite measure of the higher-order moments, *M* (see Methods). This composite measure increased significantly across days (Fig. 3C; p = 0.02).

Altogether, these analyses show that there is a consistent trend in how population activity changes across days of exposure to the same set of visual stimuli.

### Correlations Among Neurons

The sensory information encoded in the responses of a large neural population depends strongly on the nature and degree of correlation between cells (Oram et al., 1998; Sompolinsky et al., 2001). Population sparseness can be beneficial to encoding sensory information by reducing the correlation among neurons (Pitkow and Meister, 2012; Zohary et al., 1994). With these issues in mind, we calculated the correlations between the trial-averaged responses of all pairs of neurons recording in the same animal. This quantity measures the degree to which neurons are correlated due to common responses to the stimulus, also known as signal correlations (Panzeri et al., 2001; Schneidman et al., 2003). Signal correlations can only lead to redundancy between neurons, and therefore decrease the encoded information.

We plotted the histogram of all pairwise signal correlations and compared day 1 to day 6 (Fig. 4A). On day 1, neurons were strongly correlated with many pairs having correlation values above 0.7. However, on day 6, the distribution of pairwise correlations shifted to significantly lower values. The cumulative distribution of pairwise correlations showed that the decrease in correlation occurred across all values with a concentration at higher initial values (Fig. 4B). The average pairwise correlation value showed a significant decrease across all 6 days (Fig. 4C; p = 0.015).

**Figure 4.**
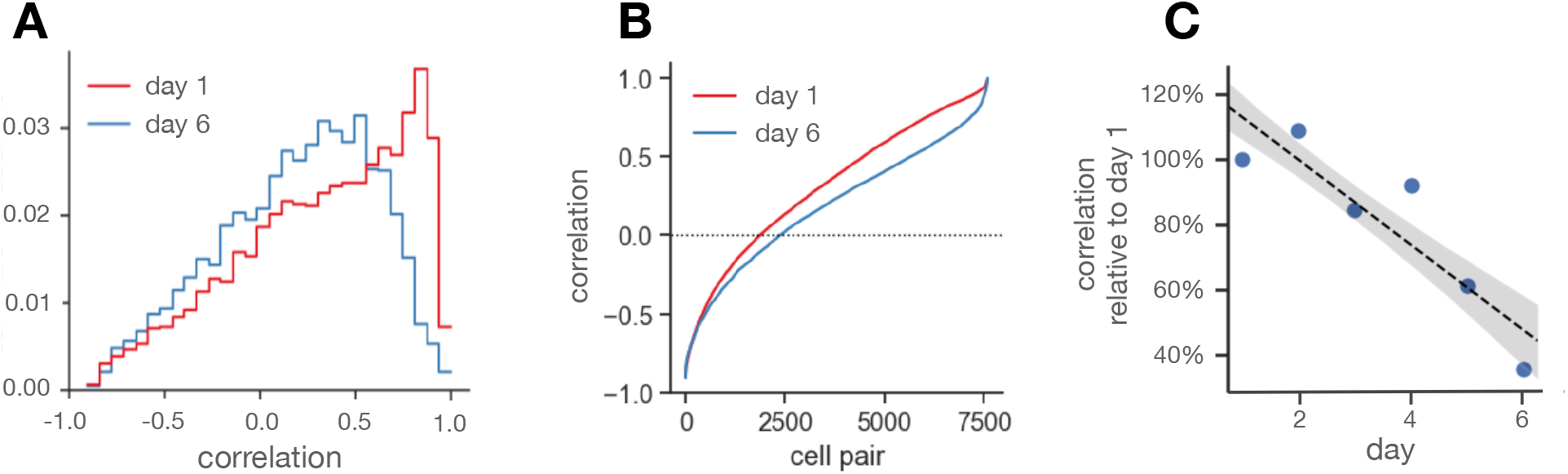
Cell to Cell Correlations. **A**. *Histogram of cell to cell correlations:* Average cell to cell corrections decreased from day 1 (red) to day 6 (blue). **B**. *Empirical cumulative distribution of cell to cell correlations*. Average cell to cell correlations decreased from day (red) 1 to day 6 (blue). **C**. *Changes in correlation across 6 days:* Correlations decreased across the 6 days of stimulus exposure (p = 0.015).

To gain further insight into the nature of changes in correlation within the population, we constructed a matrix of neuron-to-neuron correlations for each mouse. The columns and rows were sorted by the peak response times of the corresponding neurons, and the previously defined four response groups were shown by colored strips along the sides of the matrix (Fig. 5A-D). Examples from two mice showed a block diagonal structure corresponding to the four groups of neurons tuned to the same image. While the block structure was maintained across days, the overall values of correlation decreased both within and across blocks (Fig. 5 A vs. B, and C vs. D).

**Figure 5.**
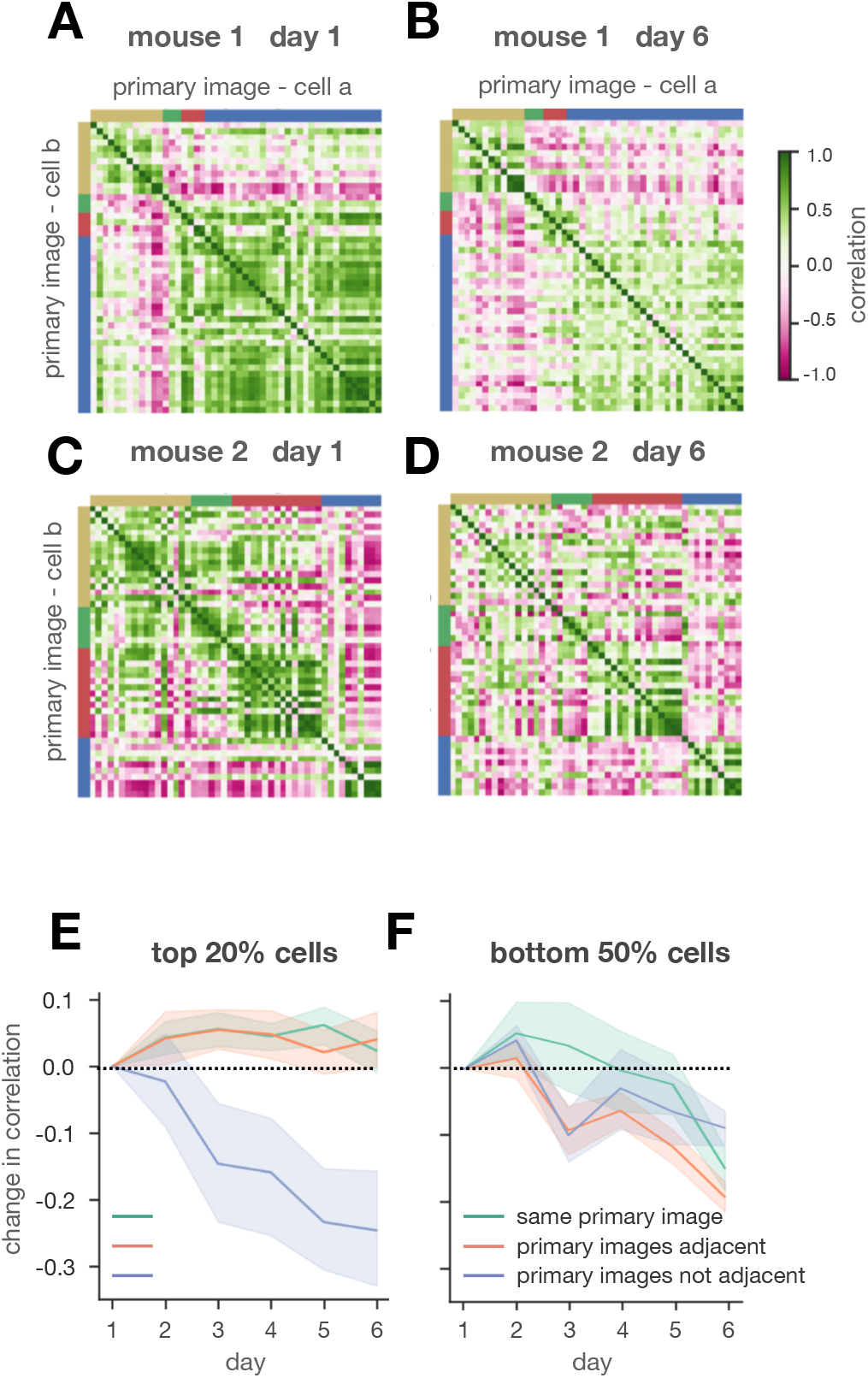
Analysis of Correlations between Neurons. **A-D**. *Correlation matrices for two example mice on day 1 versus day 6*. Panels **A** and **C** show day 1, and panels **B** and **D** show day 6. Negative correlations are red; positive correlations are green. Neurons were sorted by peak time with colors along the sides of the matrix indicating group identity (see Fig. 1F). **E, F**. *Changes in correlation across days*. Neuron pairs were categorized by image tuning: (i) both neurons from the same group (green); (ii) neurons from successive groups (red); (iii) neurons from groups separated by two images (blue). Average correlation values versus days of visual exposure were plotted for strongly responding neurons (**E**; top 20%; average correlation on day 1 = 0.065 ± 0.068) and weakly responding neurons (**F**; bottom 50%; average correlation on day 1 = 0.302 ± 0.033).

We next quantified the changes in the correlation matrix across days. In order to study changes in correlation for neurons responding to the same versus different images, we calculated the average correlation value for three conditions: (i) neurons in the same response group; (ii) neurons in response groups corresponding to successive images; (iii) neurons corresponding to response groups separated by one image. The basic logic was to compare neurons with the same tuning versus different tuning. However, because of the long timescale of calcium dynamics, category (ii) might be mixture of both. Because we found qualitatively different behaviors for strongly versus weakly responding neurons, we divided the population into neurons with the top 20% and bottom 50% of responses (Figs. 5E,F). For strongly responding neurons, the average correlation value within blocks was maintained across days (Fig. 5E, *green*). The slight increase was not statistically significant (p = 0.48). However, neurons with different image tuning showed a steady and substantial decrease in average correlation across days (Fig. 5E, *blue*). In contrast, weakly responding neurons showed a milder decrease in correlation, regardless of image tuning (Fig. 5F).

As shown above, many of the neurons developed weaker responses across six days of visual exposure (Fig. 2). Consequently, the signal-to-noise ratio of visual responses decreased across days (p = 9e-8). This leads to the possibility that decorrelation could have resulted from a relative increase in noise. To explore this possibility, we calculated average correlation for the subset of neurons that had a larger response on day 6 compared to day 1 (Supp. Fig. S2); such neurons did not show a decrease in SNR (p = 0.98). This analysis showed that average correlation stayed roughly the same for neurons with the same image tuning but decreased substantially for neurons with different image tuning (as found for strongly responding neurons in Fig. 5E). Furthermore, low-pass filtering individual ETAs did not affect these results (Supp. Fig. S2). These control analyses show that decorrelation cannot be solely explained by an increase in noise but instead results from the structure of visually driven responses.

### A Common Mechanism for Sparsification and Decorrelation

We noted previously that some neurons showed a decrease in secondary responses after multiple days of visual exposure (Fig. 2A *middle*); more such examples could be found (Fig. 6B). This observation suggests a common mechanism could underlie both sparsification and decorrelation – namely, that neurons develop more selective tuning within a set of highly exposed visual stimuli. One statistic that measures this property is *lifetime sparseness*, which has the same functional form as population sparseness but evaluates the distribution of responses across different stimuli rather than across different neurons.

**Figure 6.**
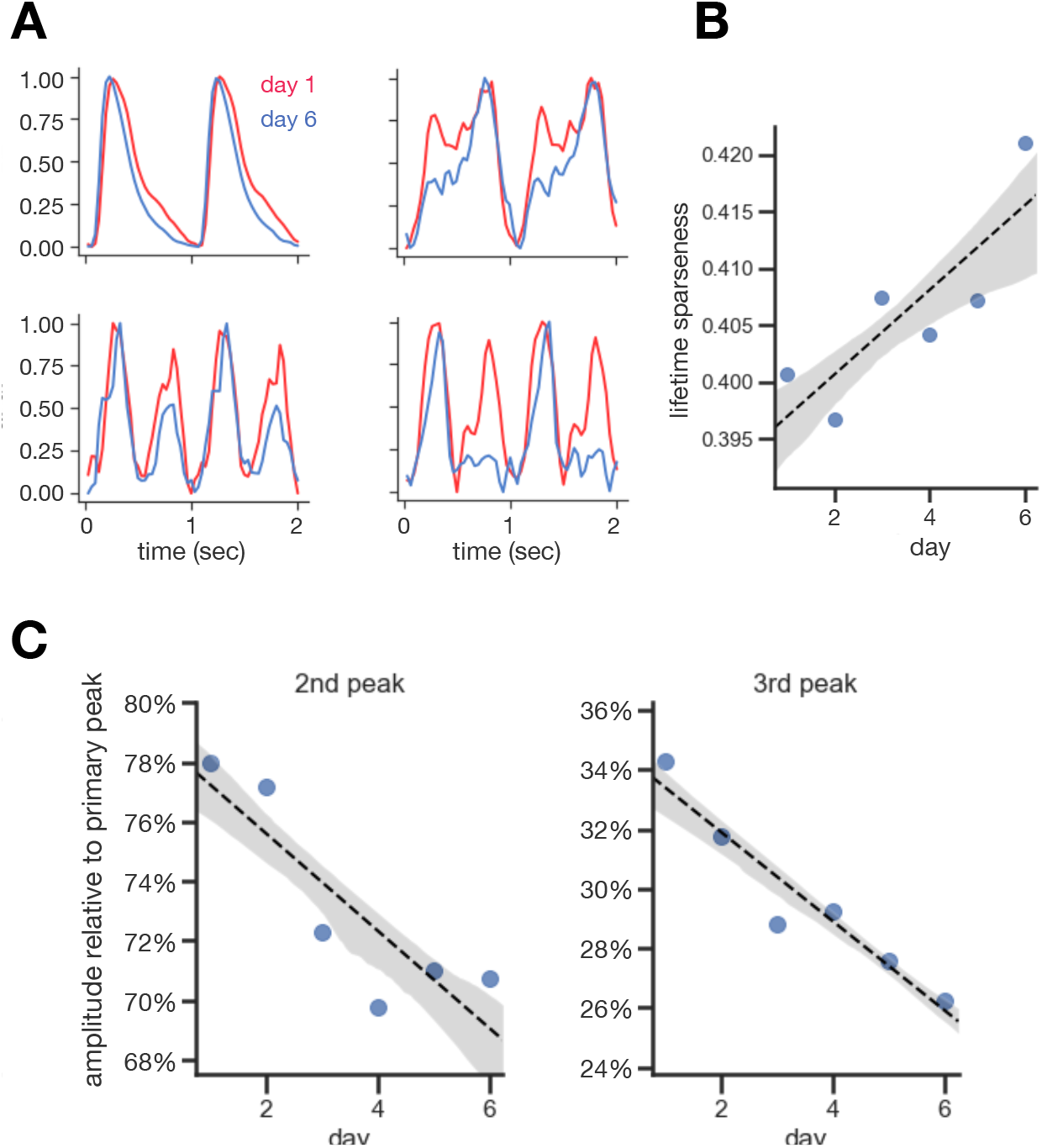
Lifetime Sparseness. **A**. *Example cells:* Red: day 1, Blue: day 6. Many strongly responding cells showed a reduction of their secondary response (if present) across 6 days of stimulus exposure. This indicates an improved tuning towards one image instead of many. **B**. *Lifetime sparseness of strongly responding cells;* Cells with a large response amplitude increased their lifetime sparseness across 6 days (p = 0.038). **C**. *Secondary responses decrease*. Averaged over cells with a mean ΔF/F > 0.2 across days, secondary and responses to additional frames decreased in amplitude compared to the primary response (2nd peak p = 0.025, 3rd peak p = 0.0027).

To determine if this effect was widespread within the population, we calculated the average lifetime sparseness of strongly responding neurons, *L*, as a function of days of visual exposure (Fig. 6A). We found a steady and significant increase across days (p = 0.015).

To further investigate, we independently rank-ordered the amplitude of each neuron’s response to the four different images (see Methods). Next, we normalized these values with respect to each neuron’s response to the primary image. This allowed us to measure how the amplitude of secondary and tertiary responses changed across days (Fig. 6C). We found that the average amplitude of secondary and tertiary responses decreased relative to the primary response. Furthermore, this effect was stronger for tertiary responses, when measured as a fractional reduction. These results give more insight into the pattern of image tuning that resulted in increased lifetime sparseness and suggests some possible circuit mechanisms (see Fig. 7).

**Figure 7.**
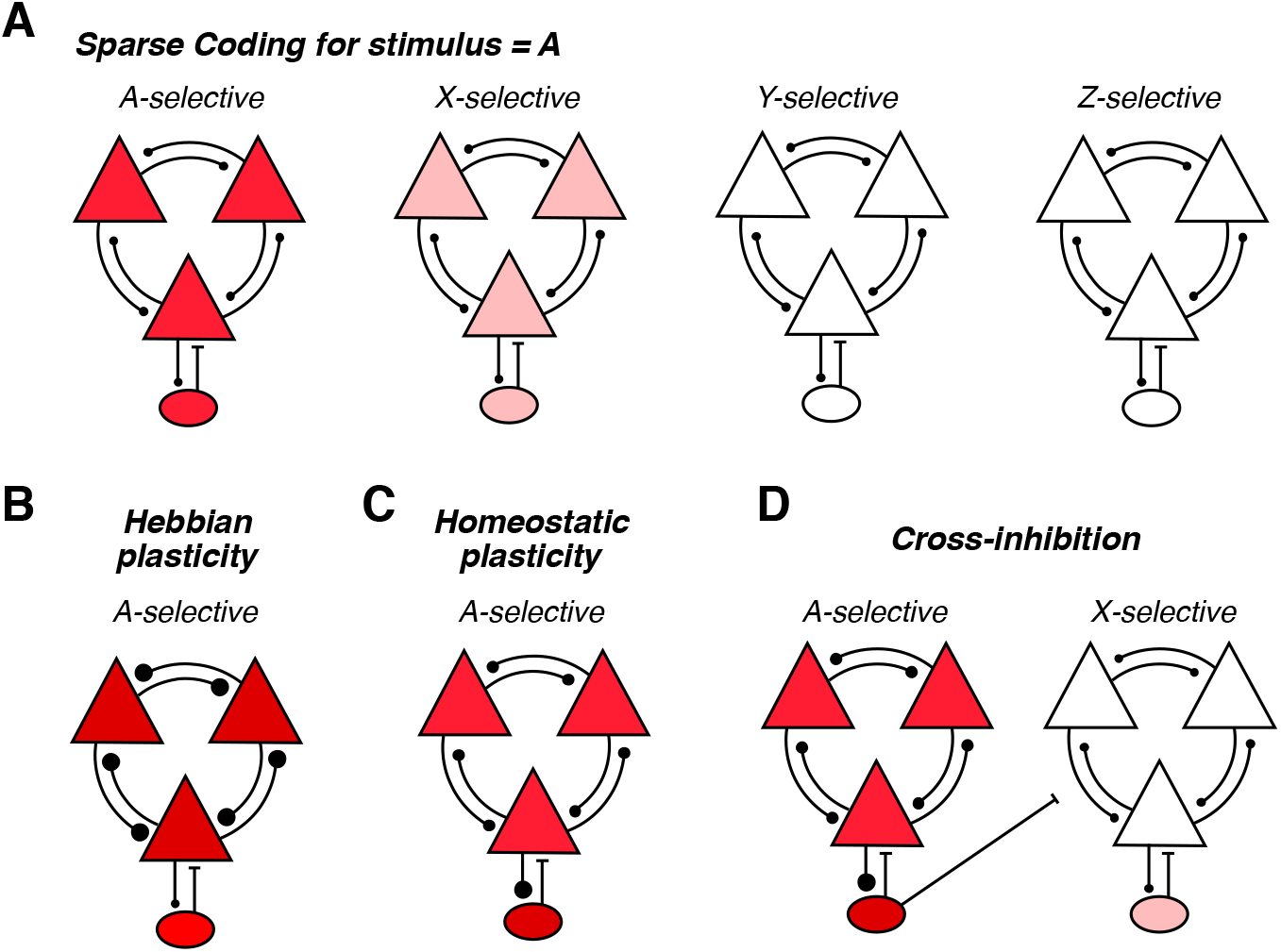
Hypothesis for the Mechanisms Underlying Sparsification. **A**. Sparse coding in a population of neurons responding to stimulus A (*triangle*: excitatory cell; *oval*: inhibitory interneuron). *Left*: A-selective neurons have high activity (red color); *Right*: neurons selective for other stimuli have low activity (pink color or no color). **B**. Upon repeated presentation of stimulus A, A-selective neurons experience Hebbian plasticity. This increases the weight of in-group synapses (larger circles) and increases activity (dark red color). **C**. Greater activity in A-selective neurons triggers homeostatic plasticity. This reduces the weight of in-group synapses and increases the weight of synapses onto inhibitory interneurons, together reducing activity to the previous level (red color). **D**. Inhibitory interneurons selective for stimulus A also make non-specific synapses onto excitatory cells selective for other stimuli. As a result of greater activity in these inhibitory interneurons, excitatory cells selective for other stimuli have their activity suppressed (no color).

## Discussion

In summary, we observed a sparsification of neural population responses to a sequence of stimuli that was repeated across several days. Weakly responsive cells decreased their responses the most, while strongly responsive cells remained relatively unchanged (Figs. 2B and 2C). Furthermore, strongly responsive cells showed increased lifetime sparseness (Fig. 6B). Signal correlations between neurons decreased across days. Weakly active cells had higher levels of initial correlation, but both weakly and strongly active cells became more decorrelated over time. For strongly active cells, decorrelation occurred primarily among neurons with different primary response frames (Fig. 5E), while for weakly active cells, decorrelation was widely distributed (Fig. 5F). Strongly responding cells decreased their sensitivity to secondary frames (Fig. 6C), which is consistent with their pattern of decorrelation. Lifetime sparseness was increased in strongly responding cells (Fig. 6B) which is also consistent with their decrease in responsiveness to secondary frames. Together, these results paint a picture in which sparsification and decorrelation are driven by a sharpening of tuning curves, together shifting population activity to represent common images more efficiently.

Many studies tracking changes in activity in the adult cortex across days with repeated exposure have involved tasks that are motivated by rewards (Henschke et al., 2020; Poort et al., 2015; Schoups et al., 2001; Woloszyn and Sheinberg, 2012). In general, these studies find that the representation of the rewarded stimulus is enhanced and the discriminability versus unrewarded stimuli is increased. However, the administration of a reward engages different plasticity mechanisms than our paradigm, including most notably plasticity at the corticostriatal synapse that is gated by dopamine (Pawlak and Kerr, 2008). Therefore, the results of these studies should be considered to be separate and potentially different from our findings for the case of passive visual exposure with no rewards. To underscore this point, a recent study found opposite effects of visual exposure depending on whether stimuli were paired with a reward or not (Henschke et al., 2020).

Importantly, passive visual exposure is a form of unsupervised learning, which can operate continuously in many different brain circuits in parallel. In contrast, dopamine elicited by rewards has a limited bandwidth. Therefore, even if plasticity due to passive exposure has a smaller impact than pairing stimuli with rewards in a specific neuroscience study, it may still have greater impact on a behaving animal due to its massive parallelism.

Some studies have explored how passive exposure to the same visual stimuli over multiple days effects neural activity in mouse V1. The average response of both L4 and L2/3 neurons to a drifting grating decreased across four days, similar to our results (Makino and Komiyama, 2015). Furthermore, the number of responsive neurons decreased across days, while the average activity of those responsive neurons remained the same. Changes were specific to the presented orientation. In another study, the number of neurons responsive to a passively repeated stimulus decreased across multiple days (Henschke et al., 2020). While there is not an exact match between what these studies called ‘responsive’ neurons and what we defined as ‘strongly’ responsive neurons, these results are broadly consistent with ours – strongly responding neurons maintained similar activity across days, while the rest of the neurons decreased their activity. This decrease could potentially result in fewer neurons meeting the definition of ‘responsive’ in both studies. One potentially relevant distinction is that Makino and Komiyama administered mild tail shocks, while we did not.

*Kim et al*. studied changes in activity of L4 neurons in V1 to the exposure of phase-reversing gratings across days and observed a decrease in average neural activity to familiar grating orientations, consistent with our findings (Kim et al., 2019b). In contrast, they observed no change in the number of responsive neurons. This study differed from ours in that its mice were not allowed to run. Kaneko et al. showed a stimulus-specific response enhancement in L2/3 neurons of V1 after several days of passive exposure to a drifting bar of a particular orientation; this enhancement resulted in sharper orientation tuning (Kaneko et al., 2017). The control group, that was not allowed to run, did not show this response enhancement. This study differed from ours in that neural responses were measured under anesthesia, which can disrupt top-down inputs to V1.

Altogether, there is some inconsistency in the effects of passive visual exposure among these studies. One possible explanation is that the results are sensitive to experimental conditions, such as running, anesthesia, or aversive conditioning. Results may also differ for neural populations in different layers of V1. Further investigation will be needed to clarify the role of these factors on passive visual learning. However, we note that among all these studies of the effects of passive visual exposure, we believe that ours used some of the most “natural” conditions: mice were allowed to run freely, neural activity was measured in the awake condition, and no tail shocks were administered.

Our results extend on these previous studies in two important ways. First, we analyzed the full distribution of population neural activity rather than simpler statistics like average activity or proportion of responsive neurons. This leads to a substantially different picture of how neural coding changes: passive visual exposure has previously been interpreted as resulting in a ‘reduced’ representation of the repeated stimulus (Henschke et al., 2020), while our results argue instead that the representation becomes more efficient. Secondly, we used multiple repeated stimuli, which allowed us to observe a decrease in signal correlations, a sharpening of tuning curves, and an increase in lifetime sparseness. Previous studies could not assess any of these properties.

What circuit mechanisms might underlie sparsification? It is now well-appreciated that multiple plasticity mechanisms constantly reshape cortical circuits in adults (Feldman, 2009; Ribic, 2020; Turrigiano, 2011). Computational models have shown that the combination of Hebbian, anti-Hebbian, and homeostatic plasticity can, in fact, increase sparseness, while decreasing correlation among neurons in a manner that preserves information (Foldiak, 1990; Pehlevan et al., 2015; Zylberberg et al., 2011). Further studies have elaborated this model (Falconbridge et al., 2006; Pehlevan et al., 2015; Zylberberg et al., 2011) and have argued that it constitutes a biologically plausible mechanism for achieving sparseness in neural circuits, as originally proposed by Olshausen and Field (Olshausen and Field, 1996a). This allows us to suggest a simple hypothesis for sparsification.

First, we start with a sparse neural code, in which a small subset of neurons in a local cortical circuit strongly responds to a given stimulus (Fig. 7A). Repeated presentation of this stimulus will lead to strengthening of synapses among responsive excitatory neurons via Hebbian plasticity (Fig. 7B). This increase in neural activity then triggers two forms of homeostatic plasticity (Fig. 7C): i) intrinsic homeostasis, which reduces synaptic strength and intrinsic excitability of the responsive neurons (Turrigiano, 2011); ii) E-I plasticity, which strengthens synapses onto inhibitory neurons that feed back onto the responsive subset of excitatory neurons (Froemke et al., 2007; Vogels et al., 2011). With the right parameters, these mechanisms could keep the activity of strongly responsive neurons roughly constant. However, the enhanced drive onto inhibitory neurons would also result in increased inhibition onto weakly responsive neurons (Fig. 7D). This would suppress the activity of the many weakly responsive neurons in the vicinity, thus leading to a sharpening of tuning curves as well as an overall reduction in average neural activity.

By keeping the activity of the few highly active neurons roughly the same, but substantially reducing the activity of the large majority of weakly active cells, the neural system can significantly reduce energy consumption without compromising the encoded information. This is because a small percentage of highly active neurons in V1 contains most of the information to decode images, while the inclusion of weakly active neurons actually can degrade performance (Yoshida and Ohki, 2020). Thus, the constancy of strong neural responses is the primary mechanism that maintains encoded information, while the suppression of weak responses may even further increase the information. Of course, for a truly optimized, cross-validated decoder, the addition of low SNR neural activity does not degrade performance (Ellis and Michaelides, 2018). But to the extent that neurons at the next stage of the sensory processing pathway cannot implement such sophisticated decoding algorithms, increased sparseness may be a simple and effective strategy to enhance encoded sensory information.

While sparsification has benefits for neural coding (Olshausen and Field, 2004), continued passive exposure to the natural environment cannot lead to perpetual sparsification, as this would eventually extinguish nearly all neural activity. Linear extrapolation of our measured rate of increase in population sparseness (Fig. 3A) would lead to the population sparseness increasing beyond one at day 25, which is not possible. This implies that the increase in sparseness will eventually taper off with further exposure. Conversely, it is logical to suggest that when stimuli cease to be repeatedly presented, sparsification of neural activity would reverse. Indeed, enhanced responses were found to revert by ∼50% after 7 days of non-exposure (Kaneko et al., 2017). Together these factors imply that the population neural code may be able to continually tune itself to the prevalence of visual stimuli encountered during natural behavior in a manner that enhances sparseness, and hence the coding efficiency, of the most common visual stimuli.

## Methods

### Animal Surgery and Husbandry

All experiments were performed according to the Guide for the Care and Use of Laboratory, and procedures were approved by Princeton University’s Animal Care and Use Committee. Mice genetically expressed GCaMP6f in excitatory neurons (Thy1 promoter, line GP5.3 Janelia Research Campus) (Chen et al., 2013b; Dana et al., 2014). Mice were implanted with a cranial window for imaging, described in detail in Dombeck et al., 2010. In short, anesthesia was induced with 2.5% isoflurane and then maintained during surgery at 1.5%. A 3 mm round hole was drilled in the skull with a dental drill in the skull at location 2 mm posterior and 1.75 lateral relative to bregma leaving the dura intact. The hole was then sealed with a cover glass attached to a metal canula. The metal ring was then glued to the bone with N-butyl cyanoacrylate (Vetbond; 3M). The skull was then covered with dental cement (Metabond; Parkell) placed in a manner that filled the space outside the ring, thus fixing the ring in place. Then, a titanium headplate was put on top of the dental cement and covered with dental cement for fixation. We allowed mice to recover for several days before recording. Pain management was provided.

### One-Photon Ca^++^ Fluorescence Imaging and Mapping of Visual Areas

In order to accurately locate V1, we mapped the visual cortex with a one-photon microscope (Garrett et al., 2014; Marshel et al., 2011). In short, awake mice were placed on an air suspended Styrofoam ball and head fixed. Drifting horizontal and vertical bars, having a flickering checkerboard pattern as their texture, were then presented to the mouse on a LCD screen. The screen covered a large part of the visual field, comprising 150 degrees vertical and 145 degrees horizontal. The bars were warped in a location-dependent fashion, in order to adjust for the viewing angle of the mouse relative to the flat surface of the monitor. While the bars were slowly moved across the visual field, the cranial window was imaged with a low magnification epi-fluorescence microscope at 30 Hz. Trials for each direction of motion were averaged and then changes in bulk activity on the surface of the brain were identified. For each identifiable visual area, a wave of activity corresponding to the bar position was observed. Those activity waves were then used to segment the visual cortex into known areas using the algorithm from (Garrett et al., 2014). For precise location identification under the 2-photon microscope, vasculature landmarks were then used.

### Two-Photon Ca^++^ Fluorescence Imaging

We imaged neural activity from mice in a custom-built imaging rig described in detail in (Dombeck et al., 2010). In brief, mice were placed in the center of a toroidal projection screen that covered the mouse’s visual field from -130 to +130 degrees horizontal and from -20 to + 70 degrees vertical. Images were projected onto this screen with a projector whose image was spread with an angular amplification mirror to cover the full toroidal screen. Images were adjusted for this distortion via software before projection. The center of the rig contained an air suspended Styrofoam ball on which mice were placed; mice were head-fixed to a post using their head plate. A two-photon microscope with a 0.88 NA, 40x water immersion objective was then used to image neural activity at cellular resolution in an area of around 400 × 400 µm at 30 Hz (512 × 512 pixel). GCamp6f was excited with a two-photon Titanium sapphire laser (140 fs pulses at 80 MHz) at 920 nm. The laser path and the data acquisition were controlled with ScanImage 5. Stimulus timing markers together with laser path mirror positions were recorded with Clampex for later synchronization. Neurons were recorded from a depth of 200 – 300 µm relative to the dura. We found the same position for fields-of-view across days using blood vessel landmarks. We compared the orientation of fields-of-view across days post-hoc; we found that these shifts in orientation were small (< 3 degrees variation).

### Design of Visual Stimuli

We formed images with a random superposition of Gabor functions (Fig. 1D). For each image, 100 Gabor functions were randomly chosen with the following properties: i) 100% contrast, ii) either ON- or OFF-polarity, iii) random location, iv) random orientation, and v) random phase. We picked the spatial extent for each Gabor function randomly from a range of 10° – 20°, as this range matches receptive field sizes in V1 neurons in mice (Niell and Stryker, 2008). Gabor functions were linearly superimposed with saturation at 100% contrast. Because of this saturation, the average light level of images varied by a small amount (standard deviation of 1.2% of the mean). Four images were presented for 250 ms each in the same temporal order and without blank frames in between for one hour each day for six days.

### Extraction of Fluorescence Traces

Due to brain motion artefacts, images had to be aligned before fluorescent trace extraction. To this end, we constructed a common reference image by averaging 1000 frames. We then shifted each individual image to achieve the highest cross correlation with the reference image. The resulting stabilized image sequence was then put through two more iterations of the same algorithm to remove any residual motion.

Fluorescence traces were extracted by averaging over pixel values for regions-of-interest (ROIs) that corresponded to individual neurons. ROIs were identified by hand using the reference image (Apthorpe et al., 2016; Homann et al., 2017). This procedure reduced activity bias. To avoid selection bias, we independently selected region of interests for each day and then only kept neurons that were identified in this manner on each of the six experimental days. This reduced the number of neurons from around 100 per mouse to ∼50 per mouse. For each ROI, the baseline fluorescence, *F*_0_(*t*), was computed for each frame (denoted by *t*) using a sliding window of 20 seconds and taking the 8th percentile of the resulting activity histogram (Domeck et al., 2010; Homann et al., 2017). Finally, we computed the fraction change in fluorescence across time, Δ*F/F*(*t*) = [*F*(*t*)– *F*_0_(*t*)]*/F*_0_(*t*).

### Response Amplitude of Neurons

We first computed the event-triggered stimulus average (ETA) of each neuron’s activity relative to the beginning of the sequence of four images, which we denote as *r*_*i*_(*t*) where *i* is the neuron index. In many figures we display this ETA over a longer range of time (2 sec); here we use a time range from 0 to 1 sec, which exactly covers one set of four images. Next, we defined the response amplitude, *R*_*i*_, as the standard deviation of the neuron’s ETA,

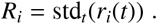

This definition robustly captured stimulus-locked changes in activity while ignoring the overall baseline activity level.

### Forming Four Groups of Neural Responses

In order to assign neurons to one of the four different response groups, we first identified for each neuron *i* the time of the peak response, *T*_*i*_, in an ETA computed over a 2 sec time window (thus including two repetitions of the set of four images). We next constructed a histogram of the peak times, *T*_*i*_. Because of the response latency, it was possible for a neuron’s primary response to occur outside of the 1 sec duration of the stimulus. In order to map all of these peak times into a one-second time interval as well as to avoid aliasing, we added into the histogram two identical sets of *T*_*i*_ that were shifted by one response cycle (±1 sec). Finally, we estimated the consensus response latency to be 133 ms. This allowed us to assign every neuron’s time of peak response into one of four groups (Fig. 1F *top*), each corresponding with the presentation of one of the images.

### Sparseness Measures

The population sparseness, *S*, was defined as:

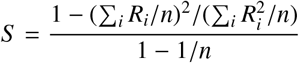

where *R*_*i*_ is the response amplitude of neuron *i* and *n* is the total number of neurons.

The inequality ratio, *I*, was defined as:

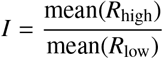

where mean(*R*_high_) is the mean of the response amplitudes of the top 10% most responsive neurons and mean(*R*_low_) is the mean of the response amplitudes of the bottom 50% least responsive neurons.

We constructed a weighted average of the higher-order moments of the distribution of log responses, *M*_*d*_ on day *d*, as follows:

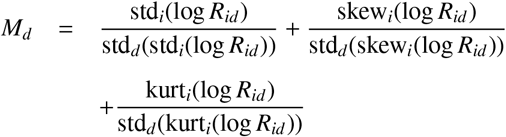

where std_*i*_(log *R*_*id*_), skew_*i*_(log *R*_*id*_) and kurt_*i*_(log *R*_*id*_) are the standard deviation, the skew and the kurtosis of the log responses *R*_*id*_ on day *d* over all neurons *i*. std_*d*_(std_*i*_(log *R*_*id*_)), std_*d*_(skew_*i*_(log *R*_*id*_)) and std_*d*_(kurt_*i*_(log *R*_*id*_)) are normalization factors to bring the three terms on a similar scale before summation. The normalization factors are equal to the standard deviation across days *d* of the corresponding statistical moments in the numerator.

The lifetime sparseness, *L*_*i*_, of a neuron *i* was defined as:

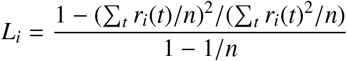

where *r*_*i*_(*t*) is the activity of neuron *i* at timepoint *t* and Σ_*t*_ denotes the sum over all timepoints and *n* is the number of timepoints that are summed over.

### Correlation Analysis

The cross correlation, *C*_*i j*_, was computed between each pair of neurons *i* and *j* recorded in the same mouse:

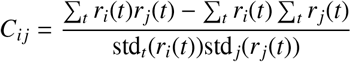

where std_*t*_(*r*_*i*_(*t*)) measures the standard deviation of a neuron’s ETA across time. This quantity (also known as the Pearson correlation coefficient) measures the signal correlation between each pair of neurons as they respond to the same set of four images.

In order to compute the correlation matrices in supplementary Figure 2, *C*_*kl*_, we calculated the average pairwise correlation among all pairs of neurons where one cells was in group *k* and the other cell was in group *l*. This operation reduced the dimensionality of the correlation matrix to 4×4. In addition, the ETA for each neuron, *r*_*i*_(*t*), was first smoothed by a Gaussian filter with a standard deviation of 66 ms.

### Non-primary Response Amplitudes

For each cell, we calculated the response amplitude to the four individual images by taking the maximum of the response traces *r*_*i*_(*t*) in the corresponding lagged response time window as defined previously (Fig. 1F). Because we were interested in relative changes across days, we normalized each neuron’s non-primary response amplitudes to the primary image on each corresponding day. This gives an amplitude of the non-primary responses relative to the primary response for each day and neuron. These normalized response amplitudes were then rank-ordered individually for each neuron. We then averaged those response amplitudes across neurons for each rank and day.

### Statistical Methods

Error bars were computed via bootstrap resampling and indicate 95% confidence intervals. Error bands in the curve fits were computed by bootstrap resampling the data 1000 times, curve fitting each resample, and then taking the envelope of the center 95% fits. This was done using the python package seaborn. In order to quote p-values in the text, we used the python statistical package statsmodels, using a t-distribution for inference.

