## Supplementary Figures for "Passive Exposure Sparsifies Neural Activity in the Primary Visual Cortex"

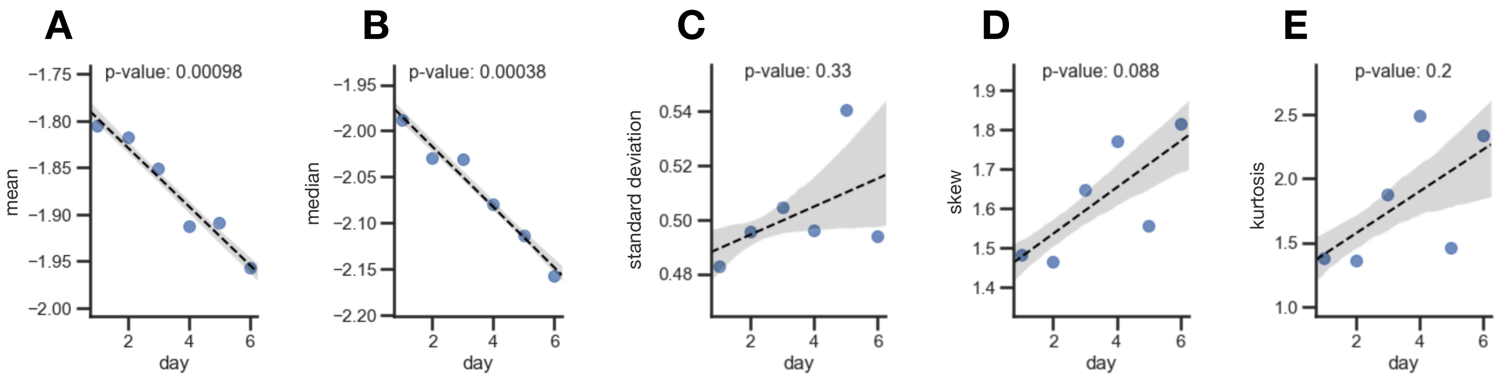


**Supplementary Figure 1: Change in Statistical Moments of the Distribution of Cell Response Amplitudes Across Days. A-E.** An average response amplitude was calculated for each cell as described in the methods section. Because the distribution of response amplitudes was very skewed (Fig. 2C) and the computed metrics are highly impacted by a few cell with large responses, we performed all measurements on log10 transformed data. Higher statistical moments tended to increase across days. **A.** *Changes in the mean of the distribution of cell response amplitudes across days.* On average cells decreased their response amplitude (p=0.00098) **B.** *Changes in median of the distribution of cell response amplitudes across days.* The median cell showed reduced activity with high confidence (p=0.00038). This implies that more than 50% of cells reduced their amplitude. **C.** *Changes in the standard deviation of the distribution of cell response amplitudes across days.* The standard deviation might have increased slightly across days (p=0.33) **D.** *Changes in the skew of the distribution of cell response amplitudes.* Skew increased across days (p=0.088) **E.** *Changes in the kurtosis of the distribution of firing rates across days.* The kurtosis of the distribution potentially increased but the scatter is large (p=0.2).

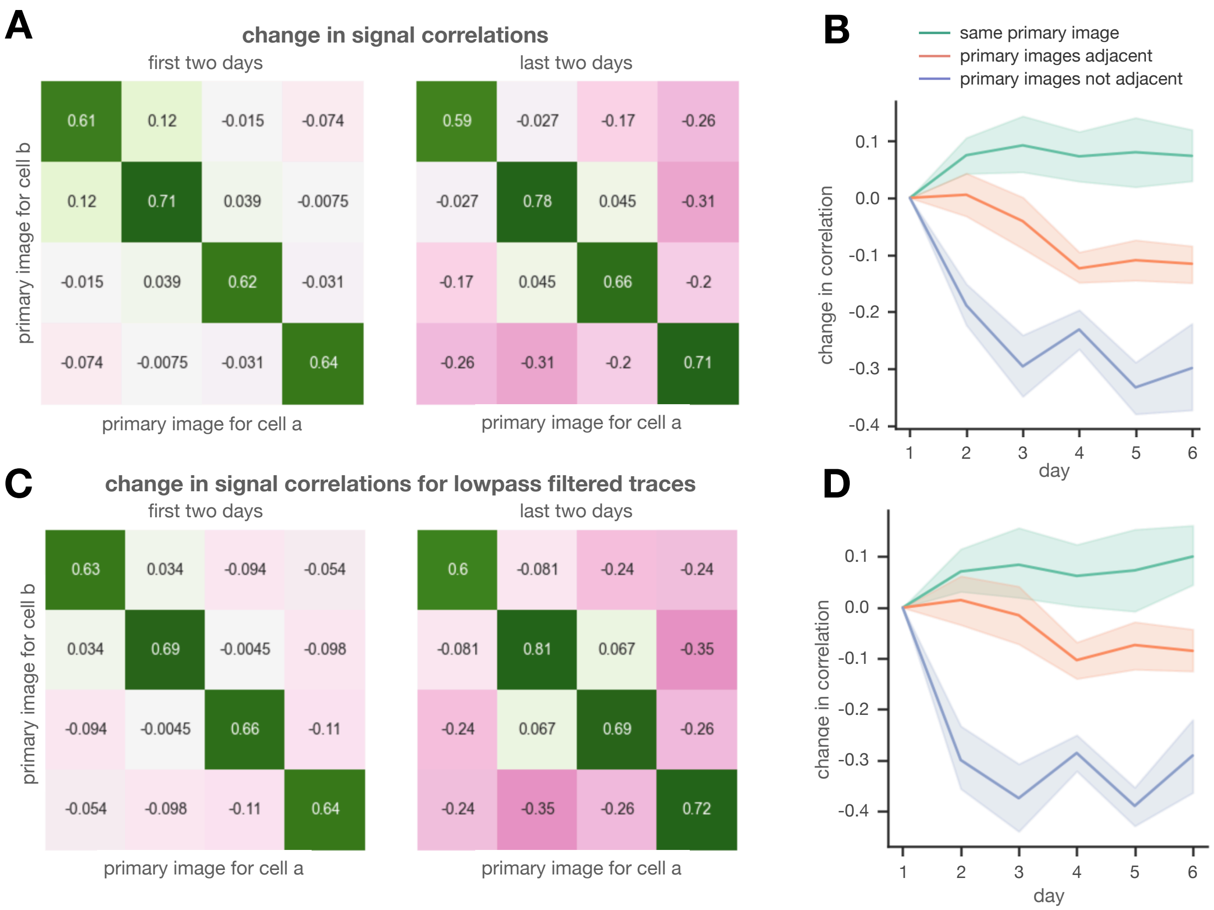


**Supplementary Figure 2: Analysis of Correlations Between Neurons with a Larger Responses on Day 6 Compared to Day 1.** Decreased correlations can potentially be caused be decreased activity due to worsening of the signal to noise ratio. We therefore investigated if cells that did not decrease in their response amplitude also decreased correlations with other cells. **A.** *Changes in signal correlation matrices across days that are block averaged for cell groups with the same primary frame.* For the left matrix, days 1 and 2 were averaged, for the right matrix, days 5 and 6 were averaged in order to improve statistical significance. Positive correlations are green, negative correlations are red. While the correlations on the diagonal increased, off-diagonal correlations decreased. **B.** *Changes in the correlation matrix across days.* Blocks were categorized by image tuning: (i) both neurons from the same group (green); (ii) blocks from successive groups (red); (iii) blocks from groups separated by two images (blue). Average correlations decreased especially for neurons that preferred images that were not consecutive. **C.** *Same as A, except that the neural responses were lowpass filtered first by a Gaussian window with a standard deviation of 66 ms.* The qualitative picture does not change, suggesting that noise is not an issue. **D.** Same as C, except for correlations derived from lowpass filtered neural traces.

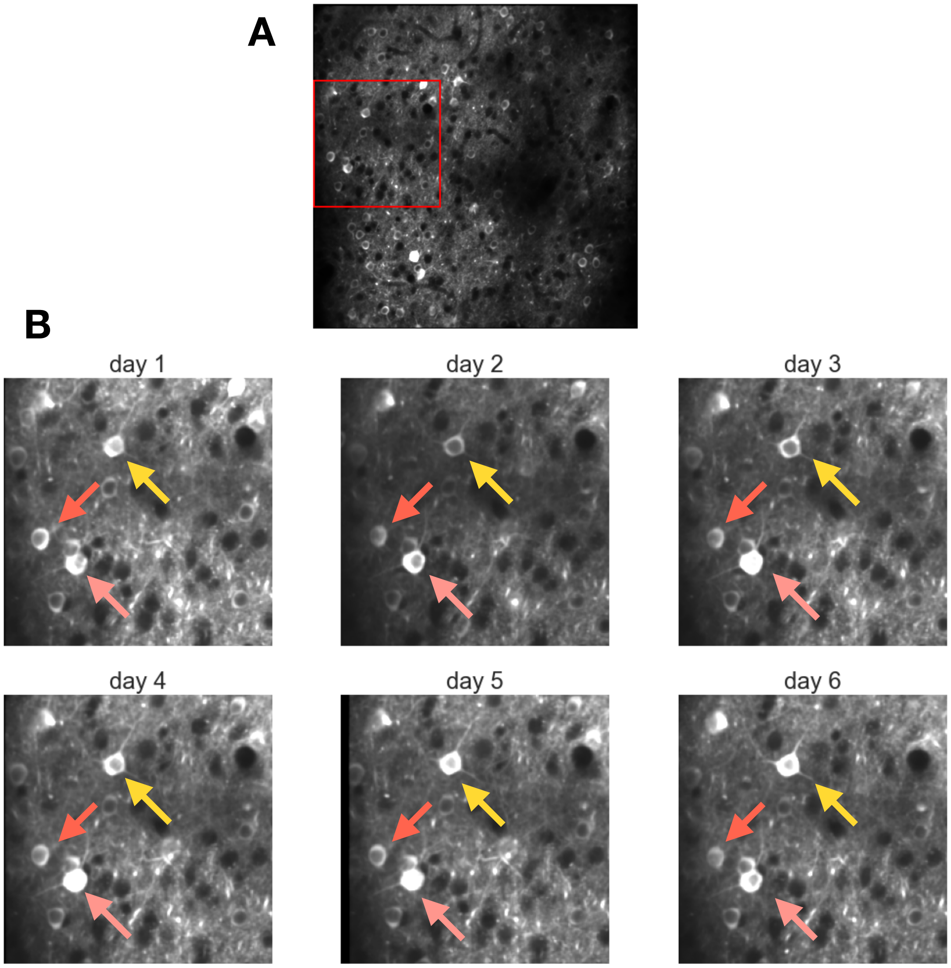


**Supplementary Figure 3: Reliable Revisiting of the Same Recording Location Across Days.** The following panels demonstrate our ability to return to the same recording location day after day to record from the same cells. Roughly 50% of neurons that were observed on at least one day could be tracked across all 6 days. **A.** *Full field of view of mouse 3 on day 1.* The region indicated by the red square is magnified in panel B. **B.** *Magnified part of the field of view across the 6 recording days.* These images show that the initial recording spot could be reliably revisited. Many neurons can clearly be seen in all 6 field of views, three of which are marked by arrows.

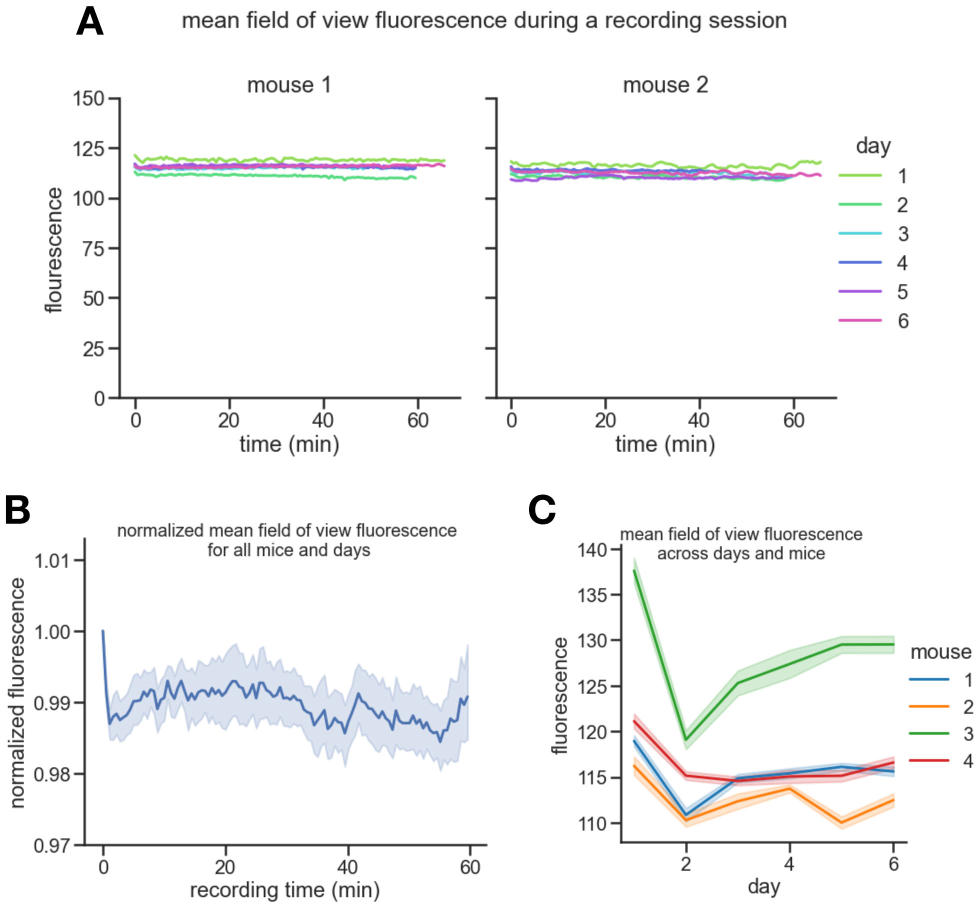


**Supplementary Figure 4: Field of View Brightness Within and Across Days.** Plots demonstrating the absence of a systematic drift in field of view brightness within and across days that could potentially create a confound in our results. **A.** *Mean fluorescence of the imaged field of view vs. recording time for two mice.* Different recording days are shown in different colors. Recording duration was roughly 60 minutes per day. Field of view brightness was stable within the 1 hour recording and across days indicating the absence of bleaching. **B.** *Mean field of view brightness averaged across all mice and days.* Traces for each mouse and day were first normalized to 1 at t=0 and then averaged. Error bands are bootstrapped 95% confidence intervals. Except for a very small drop at the beginning of the recording session, field of view brightness did not show a systematic drift during the one hour recording sessions. **C.** *Mean field of view fluorescence vs. recording day.* Different colors indicate different mice. Error bands indicate standard deviation of the field of view brightness during that particular recording session. While mean field of view brightness bounced around slightly from day to day, there was no clear trend across days.
